# Serotonin modulates social responses to stressed conspecifics via insular 5-HT_2C_ receptors in rat

**DOI:** 10.1101/2023.02.18.529065

**Authors:** Alexandra J. Ng, Lindsay K. Vincelette, Jiayi Li, Bridget H. Brady, John P. Christianson

**Author notes:** **Corresponding Author:** Alexandra J. Ng- Department of Psychology & Neuroscience, Boston College, 140 Commonwealth Ave, Chestnut Hill, MA 02467, USA.

## Abstract

Social interaction allows for the transfer of affective states among individuals, and the behaviors and expressions associated with pain and fear can evoke anxiety-like states in observers which shape subsequent social interactions. We hypothesized that social reactions to stressed individuals engage the serotonergic dorsal raphe nucleus (DRN) which promotes anxiety-like behavior via postsynaptic action of serotonin at serotonin 2C (5-HT_2C_) receptors in the forebrain. First, we inhibited the DRN by administering an agonist (8-OH-DPAT, 1µg in 0.5µL) for the inhibitory 5-HT_1A_ autoreceptors which silences 5-HT neuronal activity via G-protein coupled inward rectifying potassium channels. 8-OH-DPAT prevented the approach and avoidance, respectively, of stressed juvenile (PN30) or stressed adult (PN60) conspecifics in the social affective preference (SAP) test in rats. Similarly, systemic administration of a 5-HT_2C_ receptor antagonist (SB242084, 1mg/kg, i.p.) prevented approach and avoidance of stressed juvenile or adult conspecifics, respectively. Seeking a locus of 5-HT_2C_ action, we considered the posterior insular cortex which is critical for social affective behaviors and rich with 5-HT_2C_ receptors. SB242084 administered directly into the insular cortex (5µM in 0.5µL bilaterally) interfered with the typical approach and avoidance behaviors observed in the SAP test. Finally, using fluorescent *in situ* hybridization, we found that 5-HT_2C_ receptor mRNA (*htr2c)* is primarily colocalized with mRNA associated with excitatory glutamatergic neurons (*vglut1*) in the posterior insula. Importantly, the results of these treatments were the same in male and female rats. These data suggest that interactions with stressed others require the serotonergic DRN and that serotonin modulates social affective decision-making via action at insular 5-HT_2C_ receptors.

## 1. Introduction

When encountering others in distress, observers must assess the potential risk or reward to themselves in order to determine how to interact. Specifically, the observer must recognize and integrate environmental and social cues, evaluate personal safety, and assess the potential to help the other (Staub, 1974). In rodents, social approach and avoidance behaviors are evident in a variety of experimental paradigms allowing for direct investigation of the neurobiology underlying this type of social decision-making (Baptista-de-Souza et al., 2015; Ferretti & Papaleo, 2019; Keysers et al., 2022; S.-W. Kim et al., 2021; K. Meyza & Knapska, 2018; Rogers-Carter, Varela, et al., 2018; Sterley et al., 2018; Sterley & Bains, 2021; Walsh et al., 2023). In social interactions among rodents exposed to stress, approach or avoidance behaviors appear to be influenced by several factors including internal states, familiarity with the conspecific, nature of the stressor and the safety of the interaction context. For example, social *avoidance* occurs when rats are presented with sick conspecifics (Rieger, Worley, et al., 2022), when rats or mice are presented with stressed adult conspecifics (Ferretti et al., 2019; Rogers-Carter, Varela, et al., 2018), and when the observer is stressed themselves (Christianson et al., 2008). On the other hand, social *approach* to a stressed conspecific occurs when mice are exposed to a demonstrator expressing conditioned fear (Ferretti et al., 2019; Scheggia et al., 2020), when mice or pair-bonded voles are reunited with a stressed cagemate (Burkett et al., 2016; Sterley et al., 2018), and when rats are presented with a restrained cagemate (Bartal et al., 2011) or stressed juveniles (Rogers-Carter, Varela, et al., 2018). Further, direct observation of conspecifics in distress leads to stress-related affective behaviors and vicarious learning in the observers (For review see: (Keum & Shin, 2019; K. Z. Meyza et al., 2017; Monfils & Agee, 2019).

In these experimental paradigms, the observer rat’s behavior towards a stressed conspecific can be explained by a vicarious sense of stress or anxiety in the observer. Exposure to stressed conspecifics triggers hypothalamic pituitary adrenocortical (HPA) axis stress responses (Sterley & Bains, 2021) and increases neuronal activity in brain regions associated with emotion expression such as the amygdala, anterior cingulate, prefrontal, and insular cortices (K. Meyza & Knapska, 2018). This so-called social transfer of stress, or “emotion contagion,” is thought to be a basis for social affective and empathic responding to others (de Waal, 2008; de Waal & Preston, 2017; Keysers et al., 2022; K. Z. Meyza et al., 2017; Sterley & Bains, 2021). The clearest support of this view was reported by Sterley et al., (2018) who demonstrated that exposure to stressed conspecifics potentiated excitatory drive onto paraventricular corticotropin releasing hormone (CRH) neurons, which are critical for the initiation of the stress response. Critically, inhibiting CRH activity with a CRH antagonist or direct optogenetic inhibition prevented the observer from engaging in social interactions with the stressed conspecific. These findings reinforce the hypothesis that how an individual responds to social stress cues is strongly influenced by their own vicarious experience of stress.

Stimulation of an observer’s HPA axis by social stress cues likely leads to upregulation of complementary stress and anxiety systems in the brain. Of interest to the current study is the serotonin (5-HT) system which originates from the dorsal raphe nucleus (DRN) (Fu et al., 2010) and is activated by CRH (Kirby et al., 2008). Additionally, aversive or painful stimuli (Bouwknecht et al., 2007; Hale et al., 2008), social stressors (Abumaria et al., 2006; Wood et al., 2013), drugs of abuse (Will et al., 2004), and physiological stress (Christianson et al., 2008; Rozeske et al., 2011) all augment neural activity in the DRN 5-HT neurons. Psychological (Kawahara et al., 1993) and physical stressors (Kirby et al., 1997) increase 5-HT synthesis, extracellular 5-HT at 5-HTergic cell bodies, and postsynaptic 5-HT receptor expression (Chaouloff et al., 1999; Inoue et al., 1994). The anxiogenic effects of 5-HT are mediated by action at a large and diverse family of receptors (Hoyer et al., 2002; Waselus et al., 2011) postsynaptic to its complex network of ascending axonal projections (Waselus et al., 2006). In regards to social behavior, 5-HT action at 5-HTergic receptors in the forebrain drives both approach and avoidance behavior depending on the postsynaptic targets. Prosocial interactions are mediated by DRN projections to anterior cingulate cortex (Li et al., 2021), nucleus accumbens (Waselus et al., 2006), and oxytocin-induced release of 5-HT from the DRN (Dölen et al., 2013). In fact, optogenetic stimulation of DRN neurons and 5-HTergic projections from the DRN to the nucleus accumbens can even rescue autism-like social deficits in mice (Luo et al., 2015; Walsh et al., 2018). Additionally, social behavior modifications following uncontrollable stress and aversive social situations like social defeat are mediated by 5-HT action in the DRN, basolateral amygdala, and dorsal striatum (Christianson et al., 2010; Cooper et al., 2008; Lukkes et al., 2009; Paul et al., 2011; Strong et al., 2011a).

The anxiogenic functions of 5-HT are largely attributed to 5-HT_2C_ receptors (Drago & Serretti, 2009; Heisler et al., 2007; Marcinkiewcz et al., 2016; Millan, 2005), which are found throughout the forebrain (Clemett et al., 2000; Linley et al., 2013). For example, 5-HT_2C_ agonist administration increases the latency to investigate novel objects (Campbell & Merchant, 2003), reduces open arm exploration in the plus maze (Pockros-Burgess et al., 2014), and mimics the effects of inescapable tail shocks on later freezing (Strong et al., 2011a). 5-HT_2C_ receptor knockout mice exhibit an anxiolytic phenotype (Heisler et al., 2007) whereas 5-HT_2C_ receptor overexpression results in an anxious phenotype (Kimura et al., 2009). Consistently, 5-HT_2C_ receptor antagonists prevent anxiety-like behaviors evoked by acute 5-HT increases (Burghardt et al., 2007) and acute stress (Christianson et al., 2010) and can facilitate fear inhibition (Foilb & Christianson, 2016). Some of these actions are attributed to 5-HT_2C_ receptors in the striatum (Strong et al., 2011b) and basolateral amygdala (Christianson et al., 2010; Pockros-Burgess et al., 2014) that receive dense innervation from the DRN.

We focused on insular 5-HT_2C_ receptors in this study because the insula contributes to a range of motivated behaviors that are influenced by anxiety, including social cognition. The insula is a unique structure in social decision-making because it is necessary for both the approach and avoidance of stressed rats. Specifically, when an adult experimental rat is presented with a choice to interact with either a naive or stressed juvenile conspecific in a social affective preference (SAP) test, the adult experimental rat will spend more time interacting with the stressed juvenile. However, when presented with a pair of naive or stressed adult conspecifics, adult experimental rats avoid the stressed adult and spend more time interacting with the naive adult conspecific.

Neurobiological treatments that inhibit insular activity prevent both of these patterns (Djerdjaj et al., 2022, 2023; Rieger, Varela, et al., 2022; Rieger, Worley, et al., 2022; Rogers-Carter, Djerdjaj, et al., 2018; Rogers-Carter, Varela, et al., 2018). Because the insula contains direct input from the DRN (Gehrlach et al., 2020; Vertes, 1991; Vertes & Kocsis, 1994) and 5-HT_2C_ receptors (Clemett et al., 2000; Linley et al., 2013), we hypothesized that 5-HT and 5-HTergic input to insular 5-HT_2C_ receptors may be necessary for social affective behaviors in the SAP test.

In this study, we first tested for the necessity of the DRN 5-HT neurons in SAP tests by inhibiting the DRN with a 5-HT_1A_ agonist prior to SAP tests in male and female rats. Next, to determine if 5-HT_2C_ receptors are needed for social affective behavior, we systemically injected test rats with a 5-HT_2C_ receptor antagonist prior to SAP tests. Then, to determine if insular 5-HT_2C_ receptors specifically are needed for social affective behaviors we injected test rats with 5-HT_2C_ receptor antagonist into the posterior insula prior to SAP tests. Finally, to inform a working model for how 5-HT_2C_ receptors may impact insular physiology, we used fluorescent in situ hybridization to investigate the cellular distribution of 5-HT_2C_ receptors in the insula.

## 2. Methods

### 2.1 Animals

Male and female Sprague-Dawley rats were obtained from Charles River Laboratories (Wilmington, MA) at either age PN51 (test rats and adult conspecifics) or PN21 (juvenile conspecifics) and maintained in groups of 2-3. All animals were allowed to acclimate to the vivarium for a minimum of 7 days prior to any experimental or surgical procedures and given access to food and water *ad libitum*. Consistent with our recent experiments in female rats, on the 5 days preceding behavioral testing male and female rats were gently wrapped in a cloth towel for 1 minute to habituate the rats to experiment handling and reduce stress associated with injections (Djerdjaj et al., 2023; Rieger, Varela, et al., 2022). The vivarium maintained a 12 h light/dark cycle and all behavioral tests occurred within the first 4 h of the light cycle. All procedures and animal care were conducted in accordance with the NIH *Guide for the care and use of laboratory animals* and all experimental procedures were approved by the Boston College Institutional Animal Care and Use Committee.

### 2.2 Procedures

#### 2.2.1 Surgical implantation of indwelling cannula

While under isoflurane anesthesia (2-5% v/v in O_2_), rats were surgically implanted with bilateral guide cannula (26-gauge, Plastics One, Roanoke VA) in the insular cortex (from bregma: A/P: -1.8 mm, M/L: ± 6.5 mm, D/V: -6.8 mm from skull surface) or the DRN (from bregma: A/P: -8.1, M/L: 0, D/V: -5.1) that were affixed with stainless steel screws and acrylic dental cement. Immediately following surgery, rats were injected with analgesic meloxicam (1 mg/kg, Eloxiject, Henry Schein), antibiotic penicillin (12,000 units, Combi-pen 48, Covetrus) and Ringer’s solution (10 mL, Covetrus). Rats were then returned to their homecage and allowed at least 2 weeks (males) or 3 weeks (females) for recovery prior to behavioral testing. After behavioral testing concluded, rats were overdosed with tribromoethanol (Sigma) and brains were dissected, flash frozen, and sectioned on a cryostat at 40 µm. Slices were then mounted onto gelatin-subbed slides (Fisher) and stained with cresyl-violet to verify cannula placements by comparing microinjector tip location to the rat whole brain stereotaxic atlas. Rats were included in the analyses if their cannula were in the insula (Bregma - 1.44 through Bregma -2.8) or DRN (Bregma -7.64 through Bregma -8.3). DRN cannula confirmed to be implanted in the aqueduct were included in the analyses as the injection needle protruded 1 mm beyond the cannula tip.

#### 2.2.2 DRN 8-OH-DPAT injections

To inhibit DRN 5-HT neurons we microinjected 8-OH-DPAT, an agonist for the inhibitory 5-HT_1A_ autoreceptors which hyperpolarizes 5-HT neurons (Kirby et al., 2003) via G-protein coupled inward rectifying potassium channels (Hjorth & Magnusson, 1988). 8-OH-DPAT (Tocris) was first dissolved in 100% DMSO and then diluted in a vehicle of 10% DMSO and 0.9% saline. Injections were 1 µg in 0.5 µL and infused at a rate of 1.0 µL/min with an additional 1 min diffusion time. Testing occurred 30-40 mins after injection. This dose and preparation were selected based on prior use in our laboratory (Christianson et al., 2010) and others (Maier et al., 1995). Although we typically use a within-subjects design for pharmacology studies in the SAP test, a pilot study indicated that repeated DRN infusions were not viable so this experiment was conducted as a between-subjects design.

#### 2.2.3 Systemic and insular SB242084 injections

To determine the contribution of 5-HT_2C_ receptors we used SB242084, a brain-penetrant, competitive and selective antagonist for the 5-HT_2C_ receptor (Bromidge et al., 1997; Kennett et al., 1997). For systemic intraperitoneal (i.p.) injections, SB242084 (Tocris) was first dissolved in 100% DMSO prior to being diluted to 1 mg/kg in a vehicle of 10% DMSO and 0.9% saline. This dose was selected based on prior studies (Boulougouris et al., 2008; Burghardt et al., 2007; Christianson et al., 2010, 2013; Foilb & Christianson, 2016; Strong et al., 2011a). For insular microinjections, SB242084 was diluted to 5 µM in a vehicle of 10% DMSO and 0.9% saline, as used in prior studies (Christianson et al., 2010; Strong et al., 2011b). Injections were 0.5 µL/side at a rate of 1 µL/minute with an additional one-minute diffusion time. Systemic injections or insular infusions were made 30-40 mins prior to testing. Drug and vehicle treatments were administered in a counterbalanced, within-subjects design.

#### 2.2.4 Social Affective Preference (SAP) test

The SAP tests allow for the quantification of social interactions initiated by a test rat toward either a stressed or unstressed conspecific, providing insight into the test animal’s discrimination of socioemotional affective cues; they were conducted exactly as previously described (Djerdjaj et al., 2022, 2023; Rieger, Varela, et al., 2022; Rieger, Worley, et al., 2022; Rogers-Carter, Djerdjaj, et al., 2018; Rogers-Carter et al., 2019; Rogers-Carter, Varela, et al., 2018). Briefly, the SAP test begins when a test rat is placed in the center of an arena containing chambers for conspecifics on opposite sides of the arena. One of the conspecifics is stressed via two, 5 s 1 mA footshocks (60 s interval) immediately before testing; the other conspecific is naive to stress. The test rat is allowed to interact with the conspecifics for 5 min and time spent body sniffing and reaching for the conspecific is hand-scored by trained experimenters. In our first descriptions of the SAP test, we included analysis of a range of overt behaviors in SAP tests with male or female rats. Direct social interaction is the only objectively scorable behavior that is consistently affected by conspecific stress and age (Rogers-Carter, Djerdjaj, et al., 2018; Rogers-Carter, Varela, et al., 2018).

#### 2.2.5 RNAScope in situ fluorescent hybridization

RNAScope was performed on posterior insular cortex (Bregma -1.8) sections according to the vendor’s instructions (ACDBio). Briefly, tissue was thawed, fixed and treated with a RNAScope cocktail including probes for vesicular glutamate transporter 1 (*vglut1*, catalog #317001), glutamate decarboxylase 2 (*gad*, catalog #435801), and *htr2c* (2C, catalog #469321) mRNAs, and DAPI. The total number of nuclei (DAPI) and cells colocalized with *vglut1, gad*, and *htr2c mRNAs* were determined. Effects of sex on cell counts were assessed in two-way ANOVAs with sex as a between-group factor and hemisphere side as a within-subject factor. Data were obtained for all 3 mRNAs from both hemispheres of 4 male and 4 female rats; the *vglut1* and *htr2c* counts were replicated in a separate sample of 8 additional male and 8 female rats (both hemispheres). The mean of left and right hemisphere counts are shown.

### 2.3 Statistical Analysis

Data from experimental rats were only included if site-specificity criteria were met after verification of cannula placement. Interaction times during SAP tests were analyzed using 3-way ANOVA (conspecific age by conspecific affect by treatment) and percent preference data were analyzed using a 2-way ANOVA (conspecific age by treatment). Experiments were designed to be powerful enough to detect sex differences seeking N=12 for males and females. Rats were run in cohorts and data were analyzed sequentially after removal of subjects with misplaced cannulas resulting in some groups with a range of final sample sizes from 5 to 19. All dependent measures were initially analyzed with ANOVAs treating sex as a between subjects variable and in all cases, we found no significant main effect or interactions that included sex as a factor. This was consistent with visual inspection of the individual data which are provided in all of the data panels. (Readers interested in viewing male and female data separately are directed to Supplemental Figures 1, 2, and 3). Males and females exhibited a similar range of behavior, variance, and responses to drug treatment.

Therefore, male and female data were combined for all subsequent analysis. Main effects and interaction effects were deemed significant at p < 0.05 and followed by Sidak post hoc tests to maintain experiment-wise type 1 error rate to ɑ < 0.05. All analyses were performed in Prism (GraphPad, version 9.5.0) and results of ANOVA’s, final sample size, and post hoc test results are presented in the figure legends and in Supplemental Table 1.

## 3. Results

### 3.1 DRN microinjection of a 5-HT_1A_ agonist disrupts age and stress dependent social preference

To investigate the contributions of DRN 5-HT neurons to social emotional behaviors, we conducted SAP tests after infusion of 8-OH-DPAT or vehicle to the DRN (Figure 1). Only test rats that met site specificity requirements after cannula verification were included resulting in final sample sizes of n = (5-6) per group for females and n = (6-9) per group for males (Figure 1D). DRN administration of 8-OH-DPAT in male and female test rats prevented the approach to stressed juvenile and the avoidance of stressed adult conspecifics. There was a significant main effect of conspecific age (F_AGE_ (1, 50) = 9.436, P = 0.0034, *η*^2^= 0.074) and interaction effect of conspecific age by treatment by conspecific affect (F_AGE x DRUG x AFFECT_ (1, 50) = 45.38, P < 0.001, *η*^2^= 0.236).

**Figure 1.**
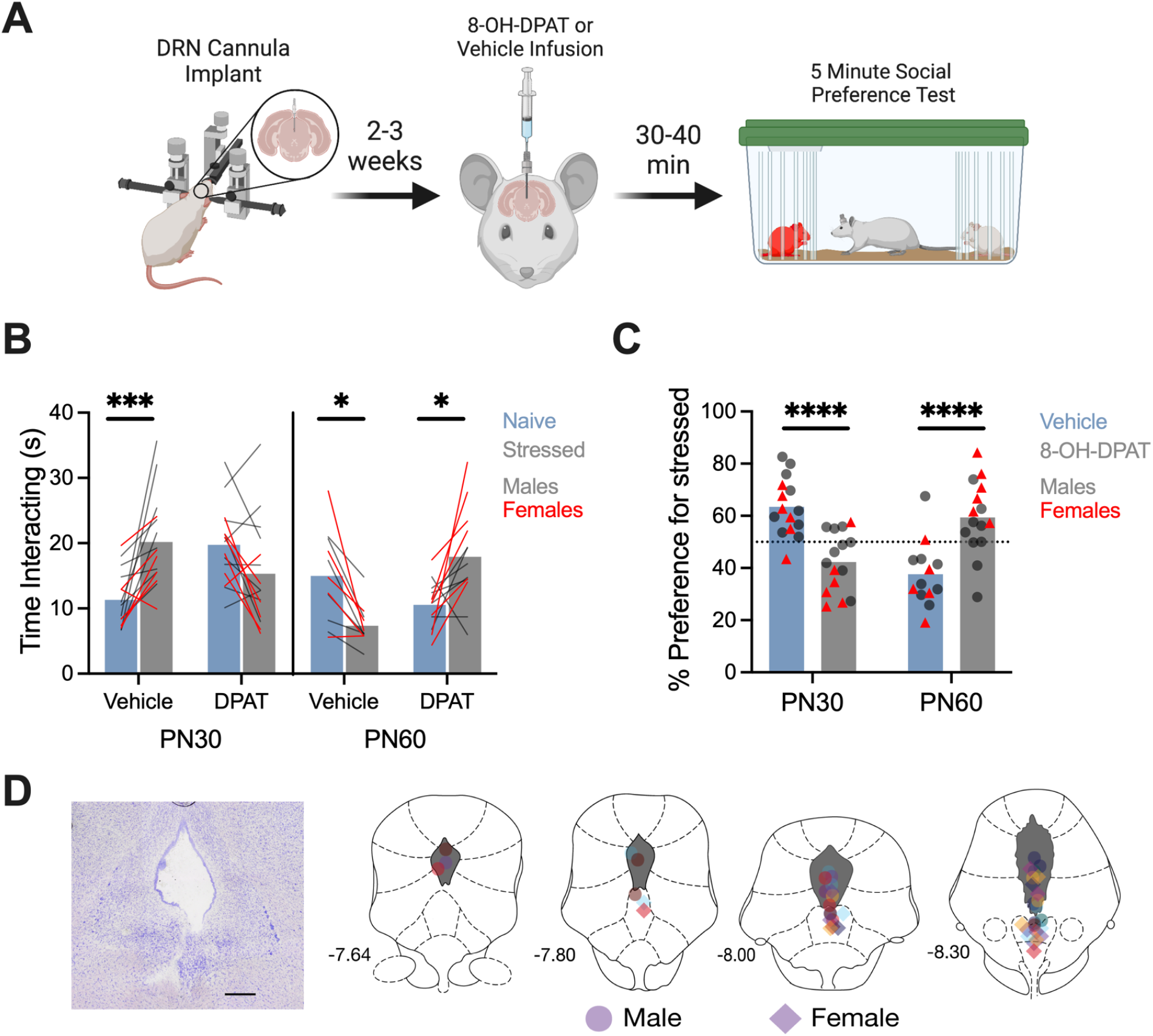
DRN microinjection of a 5-HT_1A_ agonist disrupts age and stress dependent social preference. **A**. Diagram of the social affective behavior test (SAP) paradigm. Rats received a DRN cannula implant and were given 2-3 weeks to recover. On the test day, rats received infusions of 5-HT_1A_ agonist 8-OH-DPAT (DPAT) or vehicle solution into the DRN 30-40 minutes prior to a SAP test consisting of a 5-min interaction with a naive and stressed isosexual conspecific. The amount of time spent socially investigating each conspecific was recorded. **B**. Mean (with individual replicates) time spent interacting with naive or stressed juvenile (PN30) or adult (PN60) conspecifics in the SAP test. When tested under vehicle conditions with PN30 conspecifics, male (n = 18) and female (n = 12) test rats exhibit greater exploration of the stressed conspecific (***P = 0.0005); this pattern was blocked by 5-HT_1A_ agonist 8-OH-DPAT (P = 0.2981). When tested under vehicle conditions with PN60 conspecifics, male (n = 13) and female (n = 11) test rats exhibit greater exploration of the naive conspecific (*P = 0.0191); this preference was blocked and reversed by 5-HT_1A_ agonist 8-OH-DPAT (*P = 0.0117). Male test rats are shown with black lines, and female test rats are shown with red lines. **C**. For comparison, data from B was converted to a preference score (% preference = time investigating stressed conspecific / total investigation time * 100; mean and individual replicates shown). The effect of 8-OH-DPAT was different depending on the conspecific age (F _AGE x DRUG_(1, 54) = 43.99, P < 0.001, *η*^2^= 0.443). In juvenile SAP tests, 8-OH-DPAT infusions reduced the preference for stressed conspecifics whereas in adult SAP tests, 8-OH-DPAT increased preference for stressed conspecifics (****Ps < 0.0001). **D**. A representative cresyl violet stained DRN section showing cannula damage (scale bar = 500 μm) and cannula maps showing the placement of in-dwelling DRN cannula across all experiments related to Fig. 1. Diagram in panel A was created with BioRender.com. Atlas images recreated from Paxinos and Watson (1998).

Post hoc comparison with juvenile conspecifics revealed a significant difference between the social investigation of naive and stressed juveniles in the vehicle condition (P = 0.0005) that was not present in the 8-OH-DPAT condition (P = 0.2981). Post hoc comparison with adult conspecifics revealed a significant difference between the social investigation of naive and stressed adults in the vehicle condition (P = 0.0208) and in the 8-OH-DPAT condition (P = 0.0129), in which the preference for conspecific interaction was opposite from the vehicle condition.

The preference for the stressed conspecific in each experiment was calculated as a percentage of the total time spent investigating both conspecifics (Figure 1B) and analyzed with a 2-way ANOVA. There was a significant interaction effect of conspecific age by treatment (F _AGE x DRUG_ (1, 55) = 43.99, P < 0.0001, *η*2= 0.443). Post hoc comparisons revealed a significant change in preference for stressed juveniles (P < 0.0001) and naïve adults (P < 0.0001) when comparing vehicle to 8-OH-DPAT (Figure 1C). In summary, test rats injected with vehicle preferred interaction with stressed juveniles and naive adults, while test rats injected with 8-OH-DPAT had no difference in interaction between stressed and naive juveniles and preferred interaction with stressed adults.

### 3.2 Systemic 5-HT_2C_ receptor antagonist administration interfered with social affective preference

To determine if social affective preference behavior involved the 5-HT_2C_ receptor system, we conducted SAP tests after systemic injection of the highly selective antagonist SB242084 or vehicle (Figure 2). Final group sizes were n = (8-11) for females and n = (11-18) for males. There was a significant main effect of conspecific affect (F_AFFECT_(1, 48) = 5.254, P = 0.0263, *η*^2^= 0.015), significant conspecific age by conspecific affect interaction (F_AGE x AFFECT_(1, 48) = 24.49, P < 0.0001, *η*^2^= 0.074), and a significant treatment by conspecific age by conspecific affect interaction (F_DRUG x AGE x AFFECT_(1, 48) = 47.83, P < 0.0001, *η*^2^= 0.136). Post hoc comparison with juvenile conspecifics revealed a significant difference between the social investigation of naive and stressed juveniles in the vehicle condition (P < 0.0001) that was not present in the SB242084 condition (P = 0.1709). Post hoc comparison with adult conspecifics revealed a significant difference between the social investigation of naive and stressed adults in the vehicle condition (P < 0.0001) that was not present in the SB242084 condition (P > 0.9999).

**Figure 2.**
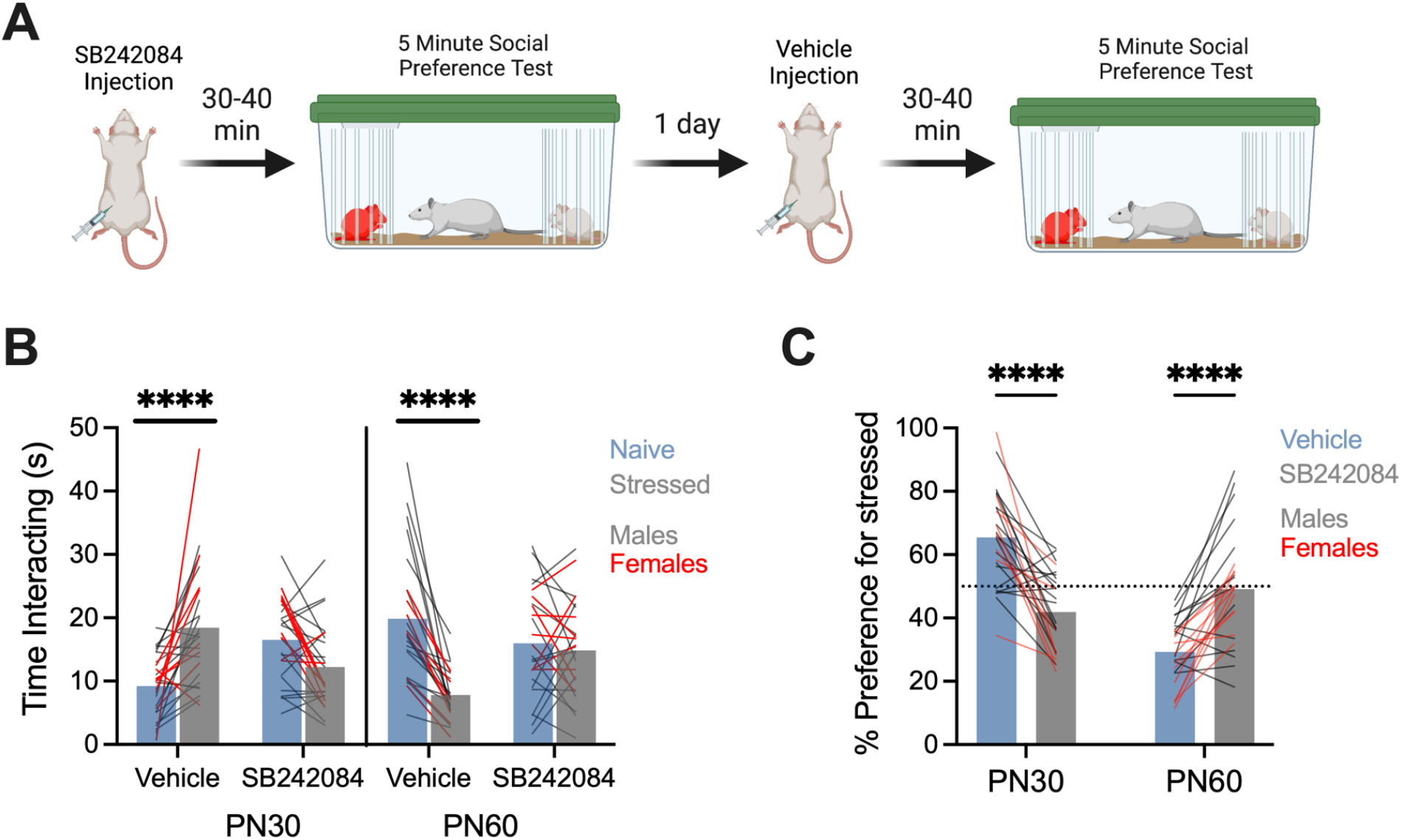
Systemic 5-HT_2C_ antagonist injections interfere with social interactions with stressed conspecifics. **A**. Diagram of the social affective behavior test (SAP) paradigm. On the test day, rats received systemic injections of 5-HT_2C_ receptor antagonist SB242084 or vehicle solution 30-40 minutes prior to a SAP test consisting of a 5-min interaction with a naive and stressed isosexual conspecific. The amount of time spent socially investigating each conspecific was recorded. **B**. Mean (with individual replicates) time spent interacting with naive or stressed juvenile (PN30) or adult (PN60) conspecifics in the SAP test. When tested under vehicle conditions with PN30 conspecifics, male (n = 18) and female (n = 8) test rats exhibit greater exploration of the stressed conspecific (****P < 0.0001); this pattern was blocked by 5-HT_2C_ antagonist SB242084. When tested under vehicle conditions with PN60 conspecifics, male (n = 11) and female (n = 11) exhibit greater exploration of the naive conspecific (P < 0.0001); this pattern was blocked by 5-HT_2C_ antagonist SB242084 (P > 0.999). Male test rats are shown with black lines, and female test rats are shown with red lines. **C**. For comparison, data from B was converted to a preference score (% preference = time investigating stressed conspecific / total investigation time * 100; mean and individual replicates shown). The effect of SB242084 was different depending on the conspecific age (F _AGE x DRUG_(1, 49) = 71.25, P < 0.0001, *η*^2^= 0.326). In juvenile SAP tests, SB242084 infusions reduced the preference for stressed conspecifics whereas in adult SAP tests, SB242084 increased preference for stressed conspecifics (****Ps < 0.0001). Diagram in panel A was created with BioRender.com.

The preference for the stressed conspecific in each experiment was calculated as a percentage of the total time spent investigating both conspecifics (Figure 2C) and analyzed with a 2-way ANOVA. There was a significant main effect of conspecific age (F_AGE_(1, 49) = 23.54, P < 0.0001, *η*^2^= 0.145) and significant conspecific age by treatment interaction (F_AGE X DRUG_(1, 49) = 71.25, P < 0.0001, *η*^2^= 0.326). Post hoc comparisons revealed a significant change in preference for stressed juveniles (P < 0.0001) and naïve adults (P < 0.0001) when comparing vehicle to SB242084 (Figure 2C). In summary, test rats injected with vehicle preferred interaction with stressed juveniles and naive adults, while test rats injected with SB242084 had no difference in interaction between stressed and naive juvenile and adult conspecifics.

### 3.3 Insular infusion of a 5-HT_2C_ receptor antagonist disrupted social affective preference

In the next experiment, we sought a locus of 5-HT_2C_ in the posterior insular cortex. SAP tests were conducted 30-40 minutes after insular infusion of SB242084 or vehicle (Figure 3A). Only test rats with cannula found to terminate in the posterior insular cortex were included resulting in final sample sizes of n = (10-11) per group for males and n = (13-18) for females. There was a significant interaction effect of treatment by conspecific age by conspecific affect (F_DRUG X AGE X AFFECT_(1, 105) = 62.31, P <0.0001, *η*^2^=15.67). Post hoc comparison with juvenile conspecifics revealed a significant difference between the social investigation of naive and stressed juveniles in the vehicle condition (P < 0.0001) that was not present in the SB242084 condition (P = 0.1798).

**Figure 3.**
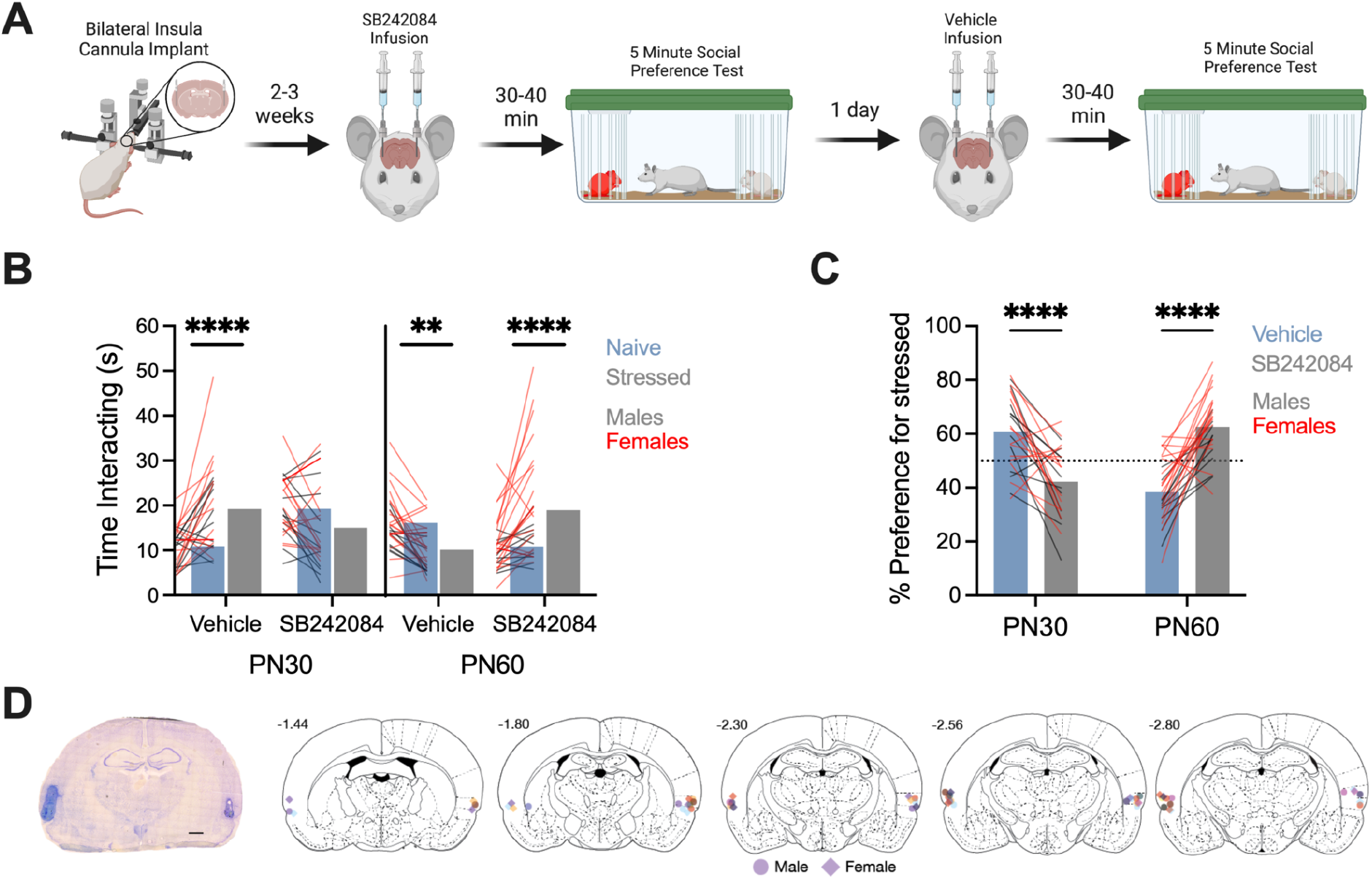
Insular 5-HT_2C_ antagonist injections interfere with social interactions with stressed conspecifics. **A**. Diagram of the social affective behavior test (SAP) paradigm. On the test day, rats received injections of 5-HT_2C_ receptor antagonist SB242084 or vehicle into the insular cortex 30-40 minutes prior to a SAP test consisting of a 5-min interaction with a naive and stressed isosexual conspecific. **B**. Mean (with individual replicates) time spent interacting with naive or stressed juvenile (PN30) or adult (PN60 conspecifics in the SAP test. When tested under vehicle with PN30 conspecifics, male (n = 11) and female (n = 14) test rats exhibit greater exploration of the stressed conspecific (****P < 0.0001); this pattern was blocked by 5-HT_2C_ antagonist SB242084. When tested under vehicle with PN60 conspecifics, male (n = 10) and female (n = 19) test rats exhibited exhibit greater exploration of the naive conspecific (**P = 0.0048); this pattern was blocked and reversed by 5-HT_2C_ antagonist SB242084 (****P < 0.0001). Male test rats are shown with black lines, and female test rats are shown with red lines. **C**. For comparison, data from B was converted to a preference score (% preference = time investigating stressed conspecific / total investigation time * 100; mean and individual replicates shown). The effect of SB242084 was different depending on the conspecific age (F_AGEx DRUG_(1, 52) = 74.8, P < 0.0001, *η*^2^=0.417). In juvenile SAP tests, SB242084 infusions reduced the preference for stressed conspecifics whereas in adult SAP tests, SB242084 increased preference for stressed conspecifics (****Ps < 0.0001). **D**. A representative cresyl violet stained IC section showing cannula damage (scale bar = 1000 μm) and cannula maps showing the placement of in-dwelling IC cannula across all experiments related to Fig. 3. Diagram in panel A was created with BioRender.com. Atlas images recreated from Paxinos and Watson (1998).

Post hoc comparison with juvenile conspecifics revealed a significant difference between the social investigation of naive and stressed adults in the vehicle condition (P = 0.0048) and in the SB242084 condition (P < 0.0001). Post hoc comparison with adult conspecifics revealed a significant difference between the social investigation of naive and stressed adults in the vehicle condition (P = 0.0048) and in the SB242084 condition (P < 0.0001), in which the preference for interaction was opposite from the vehicle condition.

The preference for the stressed conspecific in each experiment was calculated as a percentage of the total time spent investigating both conspecifics (Figure 3C) and analyzed with a 2-way ANOVA. There was a significant interaction effect of conspecific age by treatment (F _AGE x DRUG_(1, 52) = 74.8, P < 0.0001, *η*^2^=0.417). Post hoc comparisons revealed a significant change in preference for stressed juveniles (P < 0.0001) and naïve adults (P < 0.0001) when comparing vehicle to SB242084. In summary, test rats injected with vehicle preferred interaction with stressed juveniles and naive adults, while test rats injected with SB242084 had no difference in interaction between stressed and naive juvenile and a preference with interaction with stressed adult conspecifics.

### 3.4 5-HT_2C_ receptor cellular distribution and colocalization in the posterior insular cortex

To begin to understand how 5-HT action at 5-HT_2C_ receptors may impact insula function and anxiety-related behaviors, we used fluorescent *in situ* hybridization to visualize and colocalize mRNAs of 5-HT_2C_ receptor (*htr2c*) (Drago & Serretti, 2009) expressing neurons with vesicular glutamate transporter 1 (*vglut1*), a putative marker for glutamatergic neurons (Juge et al., 2006), and glutamate decarboxylase 2 (*gad*), a putative marker for GABA neurons (Ueno, 2000), in posterior insula sections from adult male and female rats (Figure 4A). Of posterior insula cells, approximately 29% were colocalized with *vglut1*, 6% with *gad* and 4% with *htr2c*. A 2-way ANOVA was performed to compare the distribution of *vglut1, gad, htr2c* mRNAs as % of total DAPI cells and sex (male vs. female). There was a main effect of mRNA type (F_mRNA_(2, 47) = 140.4, ****P < 0.001, *η*^2^= 0.8468), but no sex differences (Figure 4B). Of the cells positive for *htr2c*, approximately 70% were colocalized with *vglut1* and 20% were colocalized with *gad* (Figure 4C). A 2-way ANOVA was performed to compare the distribution of *htr2c* by cell type (*vglut1* vs. *gad*) and sex (male vs. female), which resulted in a significant main effect of mRNA type (F_mRNA_(2,31) = 55.17, ****P < 0.001, *η*^2^= 0.753) but no sex differences (Figure 4C). These results suggest that *htr2c* mRNA is concentrated in cells that express *vglut1*.

**Figure 4.**
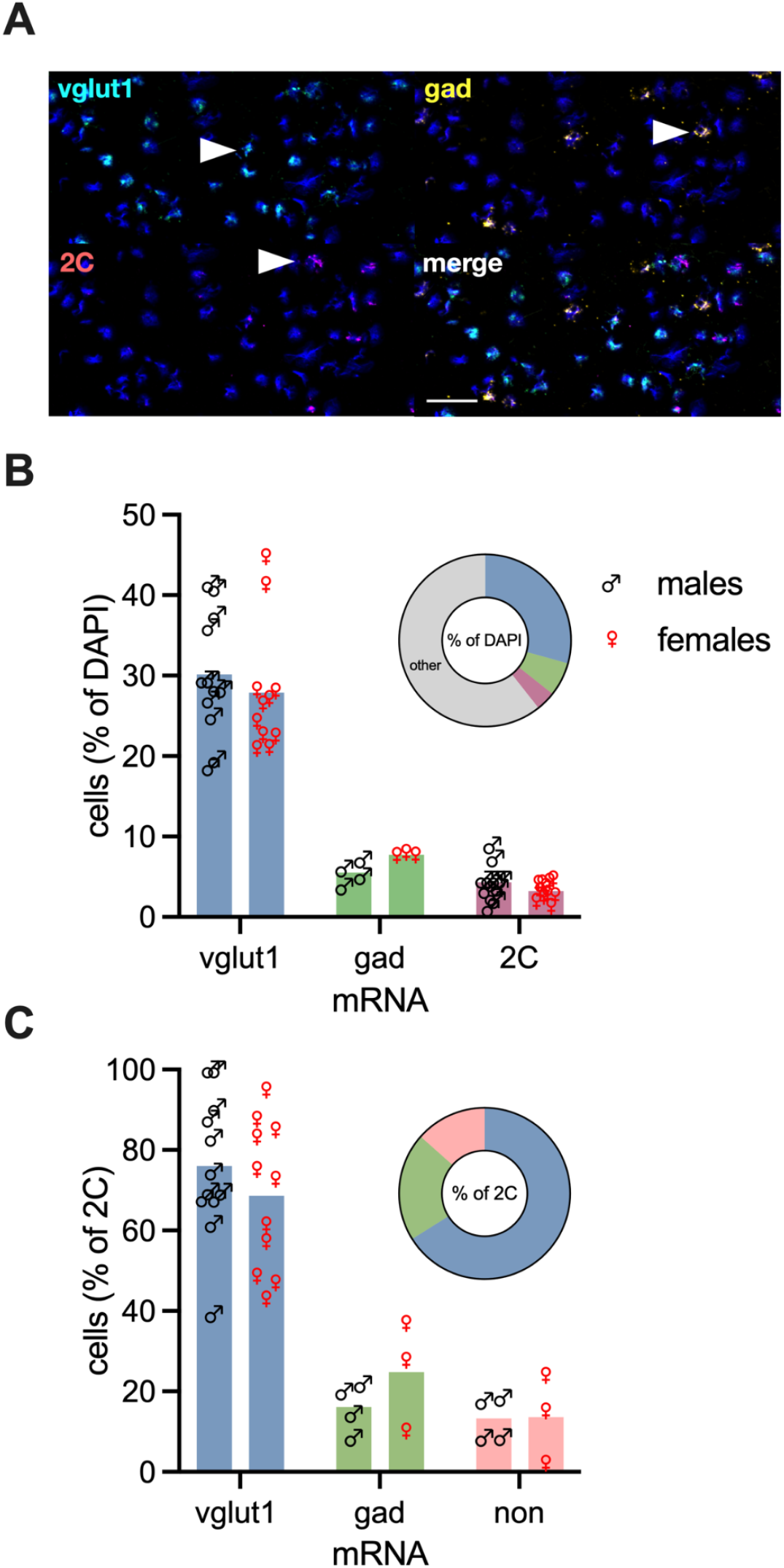
Cellular distribution of *vglut1, gad*, and *htr2c* receptor mRNA in the insular cortex. Fluorescent *in situ* hybridization was performed for *vglut1, gad, htr2c* receptor mRNAs. Fluorescent grains were counted in the left and right hemispheres of the posterior insular cortex. The total number of cells was determined by counting DAPI nuclei in each hemisphere. Nuclei containing 3 or more fluorescent grains were considered mRNA expressing cells. **A**. Representative digital photomicrographs of fluorescent visualization of DAPI and vesicular glutamate transporter 1 (*vglut1*), glutamate decarboxylase 2 (*gad*), *htr2c* (2C) mRNA from insular cortex coronal sections (20 μm), and all four channels merged. White arrows indicate representative mRNA expressing cells. Scale bar = 50 μm. **B**. Mean (with individual replicates) percent of DAPI cells colocalized with *vglut1, gad*, and *htr2c* mRNA. Data were obtained for all 3 mRNAs from 4 male and 4 female rats; the *vglut1* and *htr2c* counts were replicated in a separate sample of 8 additional male and female rats. There was a significant main effect of cell type (****F_mRNA_ (2, 47) = 140.4, P < 0.001, *η*^2^= 0.8468). There were no differences between male and female rats. The embedded pie chart shows the same data as a portion of the total DAPI cells. **C**. Mean (with individual replicates) percent of *htr2c* cells colocalized with *vglut1* and *gad* mRNA, shown as a percent of the total 2C cells per subject for male and female rats. There was a significant main effect of marker (****F_mRNA_ (2,31) = 55.17, P < 0.001, *η*^2^= 0.753). There were no differences between male and female rats. The embedded pie chart shows the same data as a portion of the total 2C cells.

## 4. Discussion

We investigated the neural mechanisms by which exposure to stressed conspecifics influences social affective decision-making. We focused on the role of 5-HT and 5-HT_2C_ receptors in the insular cortex in a SAP test in which rats typically approach stressed juveniles but avoid stressed adults. We found that inhibiting 5-HTergic neurotransmission to the forebrain via a 5-HT_1A_ agonist in the DRN and blocking systemic and insular 5-HT_2C_ receptors via a 5-HT_2C_ antagonist disrupted preference for a stressed juvenile and naive adult conspecific in the social affective preference test. *In situ* hybridization analysis revealed that 5-HT_2C_ receptor mRNA in the posterior insula are highly colocalized with excitatory glutamatergic neurons. All of these experiments included male and female rats because sex differences in the 5-HT system may exist (Carlsson & Carlsson, 1988), but there were no sex differences in any of our observations. Together, these data suggest that interactions with stressed others drives activity in the 5-HTergic DRN and that 5-HT modulates social affective decision-making via action at insular 5-HT_2C_ receptors on primarily glutamatergic neurons.

Previous studies in our lab found that neuromodulators like oxytocin (OT) and CRF are necessary during social affective decision-making in the SAP test (Rieger, Varela, et al., 2022; Rogers-Carter, Varela, et al., 2018). Although we have not tested it directly, we assume that in the SAP test, OT and CRF are released broadly through diffuse axonal projections including to the DRN where they could also affect DRN excitability, and thus, the 5-HT system overall. Relevant to social behavior, OT administration increases 5-HT binding potential in the DRN, insula, and amygdala in healthy humans, but not humans with autism spectrum disorders (Lefevre et al., 2018; Mottolese et al., 2014). CRF stimulates DRN output in higher doses (Kirby et al., 2003) to influence serotonergic neuron populations and stress-related behavior in rodents (Fox & Lowry, 2013). Related to our experiments with 5-HT_2C_ receptors, CRF enhances anxiety-related behaviors through sensitization of 5-HT_2C_ receptor signaling in cell cultures and mice (Magalhaes et al., 2010). From the perspective of the 5-HT system, 5-HT and 5-HT_2C_ receptor activation also stimulates OT release in rodents (Bagdy & Kalogeras, 1993; Jørgensen et al., 2003) suggesting that there could be interactions between social and stress related systems at multiple levels (Walsh et al., 2023). Future investigations using direct measures of these neuromodulators during behavior are needed to untangle the exact functional relationships between these systems in social behaviors.

To gain a better understanding of how 5-HT may affect insular circuit physiology we colocalized 5-HT_2C_ (htr2c) mRNA with putative markers for glutamatergic and GABAergic neurons in the posterior insula. Our results showed that about 70% of the neurons containing 5-HT_2C_ mRNA were colocalized with *vglut1*, a marker for excitatory glutamatergic neurons and ∼20% were colocalized with *gad*, a marker for GABAergic interneurons. Classically, 5-HT_2C_ receptor signals via the Gɑ_q/11_ GPCR signaling (Berg et al., 1998; Cussac et al., 2008; Gerhardt & van Heerikhuizen, 1997). Thus, 5-HT acting on 5-HT_2C_ receptors on insular glutamatergic neurons would be expected to augment local synaptic transmission and output of the insula. Similarly, OT (Zingg, 1996) and CRF (Bangasser et al., 2010) also couple to Gɑ_q/11_ and increase insular intrinsic excitability (Rieger, Varela, et al., 2022; Rogers-Carter, Varela, et al., 2018). Although a simple model in which the influx of 5-HT, OT, and CRF during social behavior augments insula synaptic efficacy and output is parsimonious, our prior work with CRF indicates that the circuitry is more complex. For example, CRF causes depolarization of insular pyramidal neurons which triggers the release of endocannabinoids which, via action of the cannabinoid type 1 receptor, suppresses presynaptic GABAergic inhibition (Rieger, Varela, et al., 2022). Another level of complexity may become evident when we can define the anatomical connectivity of insular 5-HT_2C_ expressing neurons. An interesting possibility is that certain insular projection populations may be enriched with 5-HT_2C_ receptors providing a way for social stress transfer to preferentially affect specific circuits and behaviors as recently shown in social novelty preference (S. H. Kim et al., 2022).

The current findings add to our understanding of the type of information processing and integration that occurs in the insula during social affective behaviors. Specifically, the insula integrates internal signals relating to sociability, stress, or anxiety with multimodal sensory information to augment synaptic transmission and ultimately shape the direction of behavior via outputs to prosocial or defensive subcortical systems (Rogers-Carter & Christianson, 2019). Here we speculate that the confluence of neuromodulatory signals present in the insula during social decision making may allow for greater sensory integration of multisensory inputs with motivational or affective states that is necessary to select the appropriate social behavior in a given interaction setting. The insular cortex projects to subcortical regions involved in stress and anxiety-related behaviors like the nucleus accumbens, central amygdala, and bed nucleus of the stria terminalis (Gehrlach et al., 2020; Shi & Cassell, 1998). Insula output to the NAc is necessary for prosocial approach to juveniles (Rogers-Carter et al., 2019) and 5-HT_2C_ activation may be necessary for upregulation or selecting the insular ensembles of neurons that project to the NAc. We do not yet know what insular outputs are necessary for social avoidance, however insular projections to the BNST and central nucleus of the amygdala respond to aversive stimuli and facilitate defensive behaviors (Gehrlach et al., 2019; Luchsinger et al., 2021; Schiff et al., 2018).

There are several limitations to our findings. First, we only examined the role of one 5-HT receptor found in the insula, and it is unknown whether blocking one type of 5-HTergic receptor displaces 5-HT action to other 5-HTergic receptors. Since the insula is rich with 5-HT_1A_, 5-HT_2A_, and 5-HT_2C_ receptors (Clemett et al., 2000; Ju et al., 2020; Linley et al., 2013), it is possible that 5-HT_1A_ and/or 5-HT_2A_ receptors are preferentially activated when 5-HT_2C_ receptors are blocked.

This is relevant to our experiments of social stress signals since the binding potentials of 5-HT_1A_ and 5-HT_2A_ receptors in the insula are correlated with anxiety in humans and non-human primates (Akimova et al., 2009), and 5-HT_1A_ receptor binding in the insula and DRN is lower in humans with social anxiety disorder (Lanzenberger et al., 2007). Regarding cellular distribution, insular multisensory integration is associated with parvalbumin GABAergic interneurons (Gogolla et al., 2014). Our results reveal that about 20% of the *htr2c* mRNA colocalized with a general marker of GABAergic neurons; it would be interesting to determine if specific interneuron subtypes are enriched with 5-HT_2C_ receptors. Finally, we had no direct measure of 5-HT release in the DRN or insula during the SAP test. Thus, important goals of future work include determining potential displacement of 5-HT action among different insular 5-HTergic receptors, the cellular mechanism by which social stress signals activate the DRN, if 5-HTergic fibers from the DRN to the insula are involved in social affective behaviors, and how different neuromodulators in the insula interact and affect synaptic excitability to influence social affective behaviors.

The insula is implicated in autonomic control, decision-making, self-awareness, social cognition, and integration of interoceptive and exteroceptive cues (Gogolla, 2017; Paulus & Stein, 2006; Rogers-Carter & Christianson, 2019). In anxiety-related disorders like PTSD and social anxiety, altered interoception and heightened sense of self-danger leads to an anxiety bias, which causes individuals to behave as if safe or ambiguous stimuli are relatively more threatening.

Importantly there are noteworthy sex differences in the prevalence of these disorders, sex differences in rodent social behavior (Dumais & Veenema, 2016) and sex-specific effects of CRF and OT in the insula of the SAP test (Djerdjaj et al., 2023; Rieger, Varela, et al., 2022; Rogers-Carter, Varela, et al., 2018). Although prior reports note some sex differences in 5-HT immunoreactivity and neurochemistry (Carlsson & Carlsson, 1988), we did not find any evidence of sex differences in our behavioral, pharmacological or mRNA colocalization experiments. The current findings suggest that 5-HT contributes to insular and broader decision-making networks in response to exposure to social stress signals and encourages further investigation of insular 5-HT action in anxiety and stress-related behavioral responses.

## Author Contributions

Conceptualization: AJN, LKV, JPC; methodology: AJN, LKV, JPC; investigation: AJN, JL, BHB, LKV, JPC; writing (original draft): AJN, JPC; writing (revision and editing): AJN, JPC; funding acquisition: JPC.

## Acknowledgements

The authors wish to thank Nancy McGilloway and Todd Gaines, administrators of the Boston College Animal Care Facility, for outstanding animal husbandry, Bret Judson, director of the Boston College Imaging Core, for training and assistance with microscopy, Alexis Callen for assistance with behavioral testing, and Sara Barry and Shemar Joseph for assistance with RNAScope. Funding for this work was provided by Boston College Undergraduate Research Fellowship and the National Institutes of Health Grant MH119422 to JPC.

