## Supplemental Figures for "Serotonin modulates social responses to stressed conspecifics via insular 5-HT_2C_ receptors in rat"

Supplemental Table 1. Summary of Statistics

| Panel | Effect | F Statistic | P value | $\eta 2$ |
|---|---|---|---|---|
| **Figure 1B: DRN 8-0H-DPAT SAP's** | | | | |
| | Conspecific Age | F (1, 50) = 9.436 | **P=0.0034** | 0.07386 |
| | Treatment | F (1, 50) = 3.559 | P=0.0651 | 0.02785 |
| | Conspecific Affect | F (1, 50) = 0.9760 | P=0.3280 | 0.005081 |
| | Conspecific Age x Treatment | F (1, 50) = 0.2438 | P=0.6236 | 0.001909 |
| | Conspecific Age x Conspecific Affect | F (1, 50) = 1.241 | P=0.2706 | 0.00646 |
| | Treatment x Conspecific Affect | F (1, 50) = 0.1641 | P=0.6871 | 0.0008544 |
| | Conspecific Age x Treatment x Conspecific Affect | F (1, 50) = 45.38 | **P<0.0001** | 0.2362 |
| **Figure 1C: DRN 8-0H-DPAT % Preference** | | | | |
| | Interaction | F (1, 54) = 43.99 | **P<0.0001** | 0.4432 |
| | Conspecfic Age | F (1, 54) = 1.843 | P=0.1803 | 0.01856 |
| | Treatment | F (1, 54) = 0.007990 | P=0.9291 | 0.00008049 |
| **Figure 2B: Systemic SB242084 SAP's** | | | | |
| | Treatment | F (1, 48) = 0.9325 | P=0.3390 | 0.003981 |
| | Conspecific Age | F (1, 48) = 0.1859 | P=0.6683 | 0.001089 |
| | Conspecific Affect | F (1, 48) = 5.254 | **P=0.0263** | 0.01587 |
| | Treatment x Conspecific Age | F (1, 48) = 0.2390 | P=0.6271 | 0.00102 |
| | Treatment x Conspecific Affect | F (1, 48) = 0.5226 | P=0.4733 | 0.001488 |
| | Conspecific Age x Conspecific Affect | F (1, 48) = 24.49 | **P<0.0001** | 0.07397 |
| | Treatment x Conspecific Age x Conspecific Affect | F (1, 48) = 47.83 | **P<0.0001** | 0.1362 |
| **Figure 2C: Systemic SB242084 % Preference** | | | | |
| | Conspecific Age x Treatment | F (1, 49) = 71.25 | **P<0.0001** | 0.326 |
| | Conspecific Age | F (1, 49) = 23.54 | **P<0.0001** | 0.1448 |
| | Treatment | F (1, 49) = 0.5577 | P=0.4587 | 0.002552 |
| | Rat | F (49, 49) = 1.344 | P=0.1520 | 0.3013 |
| **Figure 3B: Insular SB242084 SAP's** | | | | |
| | Treatment | F (1, 105) = 2.624 | P=0.1083 | 0.01346 |
| | Conspecific Age | F (1, 105) = 2.970 | P=0.0878 | 0.01523 |
| | Conspecific Affect | F (1, 105) = 3.329 | P=0.0709 | 0.008372 |
| | Treatment x Conspecific Age | F (1, 105) = 0.01269 | P=0.9105 | 0.00006507 |
| | Treatment x Conspecific Affect | F (1, 105) = 0.1878 | P=0.6656 | 0.0004724 |
| | Conspecific Age x Conspecific Affect | F (1, 105) = 0.3129 | P=0.5771 | 0.0007871 |
| | Treatment x Conspecific Age x Conspecific Affect | F (1, 105) = 62.31 | **P<0.0001** | 0.1567 |
| **Figure 3C: Insular SB242084 % Preference** | | | | |
| | Conspecific Age x Treatment | F (1, 52) = 74.80 | **P<0.0001** | 0.4177 |
| | Conspecific Age | F (1, 52) = 0.1949 | P=0.6607 | 0.001027 |
| | Treatment | F (1, 52) = 1.204 | P=0.2776 | 0.006722 |
| | Rat | F (52, 52) = 0.9436 | P=0.5825 | 0.274 |
| **Figure 4B: RNAScope markers as %DAPI** | | | | |
| | Interaction | F (2, 47) = 0.4708 | P=0.6274 | 0.00284 |
| | Marker | F (2, 47) = 140.4 | **P<0.0001** | 0.8468 |
| | Sex | F (1, 47) = 0.04520 | P=0.8326 | 0.0001363 |
| **Figure 4C: RNAScope markers as %2C** | | | | |
| | Interaction | F (2, 31) = 0.7573 | P=0.4774 | 0.01034 |
| | Marker | F (2, 31) = 55.17 | **P<0.0001** | 0.753 |
| | Sex | F (1, 31) = 0.007716 | P=0.9306 | 0.00005266 |

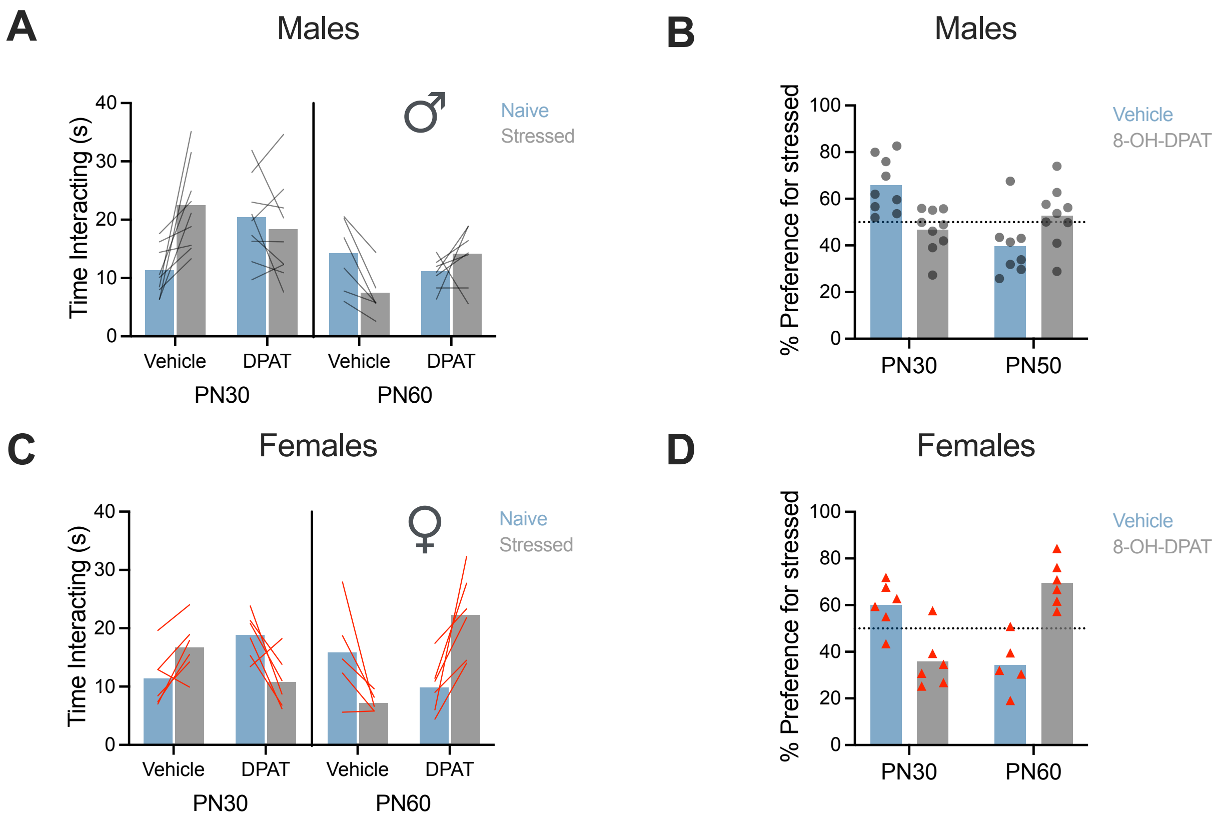


**Supplemental Figure 1. DRN microinjection of a 5-$HT_{1A}$ agonist (1 µg in 0.5 µL) disrupts age and stress dependent social preference. Data from Figure 1 separated by sex. A.** Mean (with individual replicates) time spent interacting with naive or stressed juvenile (PN30) or adult (PN60) conspecifics in the SAP test for males only. **B.** For comparison, data from A was converted to a preference score (% preference = time investigating stressed conspecific / total investigation time * 100; mean and individual replicates shown) for males only. **C.** Mean (with individual replicates) time spent interacting with naive or stressed juvenile (PN30) or adult (PN60) conspecifics in the SAP test for females only. **D.** For comparison, data from C was converted to a preference score (% preference = time investigating stressed conspecific / total investigation time * 100; mean and individual replicates shown) for females only. See main text for statistical results.

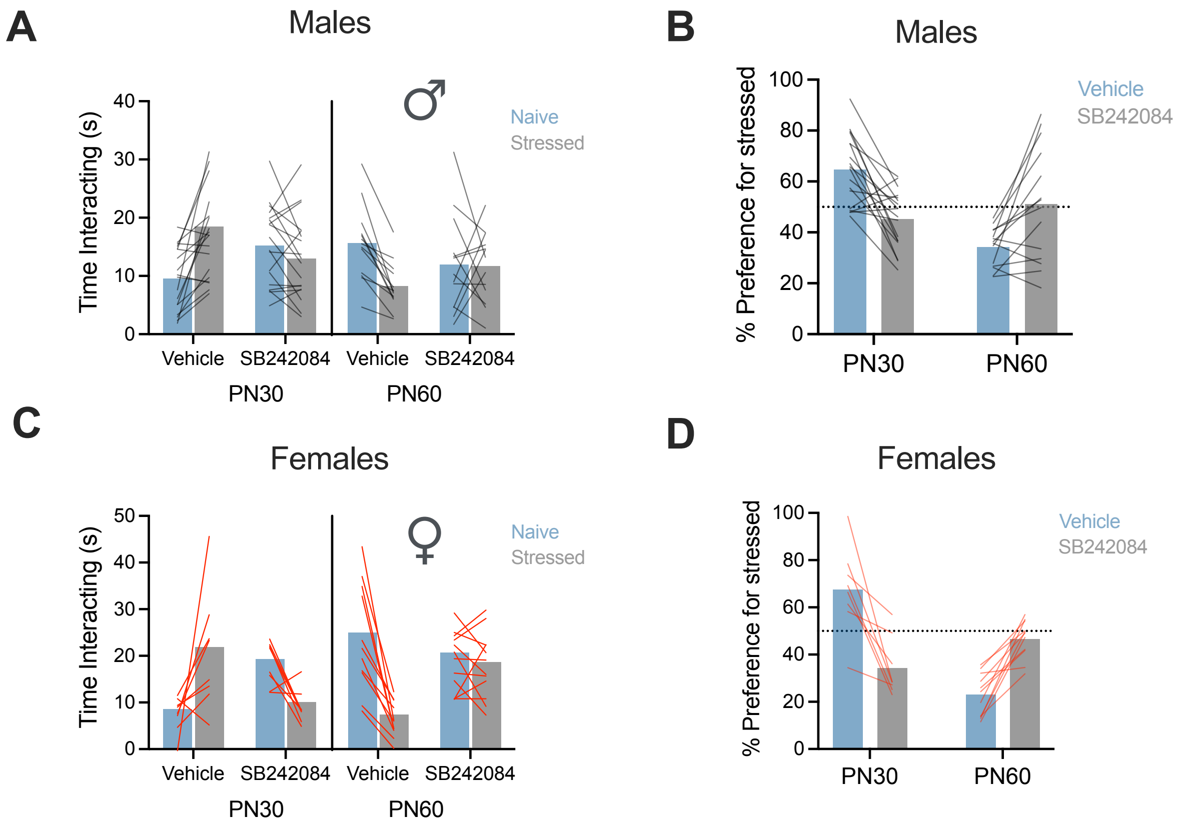


**Supplemental Figure 2. Systemic administration of 5-HT$_{2C}$ antagonist SB242084 (1mg/kg) interferes with age and stress dependent social preferences. Data from Figure 2 separated by sex. A.** Mean (with individual replicates) time spent interacting with naive or stressed juvenile (PN30) or adult (PN60) conspecifics in the SAP test for males only. **B.** For comparison, data from A was converted to a preference score (% preference = time investigating stressed conspecific / total investigation time * 100; mean and individual replicates shown) for males only. **C.** Mean (with individual replicates) time spent interacting with naive or stressed juvenile (PN30) or adult (PN50) conspecifics in the SAP test for females only. **D.** For comparison, data from C was converted to a preference score (% preference = time investigating stressed conspecific / total investigation time * 100; mean and individual replicates shown) for females only. See main text for statistical results.

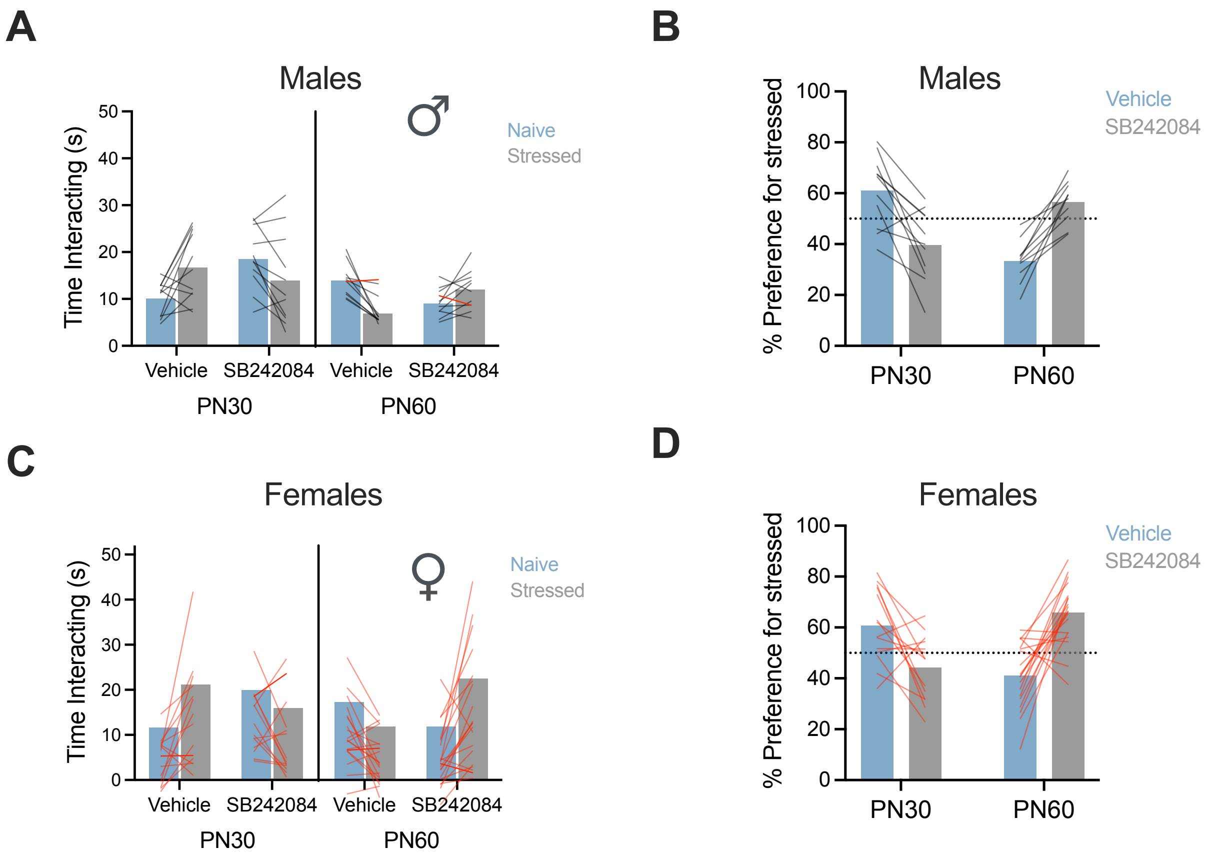


**Supplemental Figure 3. Insular administration of 5-HT2C antagonist SB242084 (5uM) interferes with age and stress dependent social preferences. Data from Figure 3 separated by sex. A.** Mean (with individual replicates) time spent interacting with naive or stressed juvenile (PN30) or adult (PN60) conspecifics in the SAP test for males only. **B.** For comparison, data from A was converted to a preference score (% preference = time investigating stressed conspecific / total investigation time * 100; mean and individual replicates shown) for males only. **C.** Mean (with individual replicates) time spent interacting with naive or stressed juvenile (PN30) or adult (PN60) conspecifics in the SAP test for females only. **D.** For comparison, data from C was converted to a preference score (% preference = time investigating stressed conspecific / total investigation time * 100; mean and individual replicates shown) for females only. See main text for statistical results.
